# Sustained release and pharmacologic effects of human glucagon-like peptide-1 and liraglutide from polymeric microparticles

**DOI:** 10.1101/262782

**Authors:** Luis Peña Icart, Fernando Gomes de Souza, Luís Maurício T. R. Lima

## Abstract

The GLP-1 class of peptide agonists has been shown to exert regulatory key roles in both diabetes, obesity and related complications. Given the short half-life of GLP-1 its use has been historically discouraged. We developed polymeric microparticles loaded with either human GLP-1 (7-37) or liraglutide peptides by double emulsion and solvent evaporation approach. The size distribution of all formulations was of about 30-50 μm. The *in vitro* kinetic release assays showed a sustained release of the peptides extending up to 30 to 40 days with varying profiles. Morphologic analysis demonstrated a more regular particle surface for those comprising polymers PLA, PLA-PEG and PLGA. *In vivo* evaluation in Swiss male mice demonstrated a similar extension of effect of decreasing in body weight gain for up to 25 days after a single subcutaneous administration of either hGLP-1 or liraglutide peptide-loaded microparticles (200 μg peptide / kg body weight) compared to controls. These demonstrate the effectiveness of hGLP-1 as a therapeutic agent in long-term, continuous release from peptide-load microparticles, and thus its plausibility as an unmodified therapeutic agent.

## 1 Introduction

The glucagon-like peptide-1 (GLP-1) is a natural peptide secreted in response to meal that stimulates glucose-dependent insulin release and suppresses postprandial glucagon secretion. Also this hormone delays gastric emptying, helping to increase satiety (Zander et al., 2002). The GLP-1 is rapidly metabolized by dypeptidil-peptidide IV (DPP4) enzyme, resulting in a half-life of about 10 min (Holst, 2007).

The physiological and pharmacological features of GLP-1 is highly attractive, in particular for additional cardiovascular outcomes and favorable risk/benefit balance, having motivated the development of several agonists and prospection of analogues (Bethel et al., 2018; Kim and Kim, 2017; Marso et al., 2016). Currently two main products are available commercially for therapeutic purposes, known as liraglutide and exenatide (exendin-4). Liraglutide is a GLP-1 analogue to which the amino acid substitution of lysine with arginine at position 34 and attachment of a C16 acyl chain via a glutamoyl spacer to lysine at position 26, resulting in a compound with increased half-life (Agersø and Vicini, 2003), decreased susceptibility to digestion by DPP4 (Ahrén and Schmitz, 2004; Brunton, 2014), and is currently available as injectable solution and allowing a s.i.d. therapeutic scheme (Gough, 2012; Ponzani, 2013; Sjöholm, 2010) either alone (Victoza^®^) or in combination with long-acting insulin analogue degludec (Tresiba).

Exending is a GLP-1 peptide analogue discovered from the salive of guila monster, which display GLP-1 agonism pharmacologic effect, and currently marketed as a pos-prandial injection or a once-a-week or once-a-month formulation in polymeric microparticles (Liu et al., 2010; Minze et al., 2013; Pinelli and Hurren, 2011). An adverse reaction observed with exendin therapy is pancreatite (Aroda and DeYoung, 2011; Ryan et al., 2013a), and without the cardiovascular benefits observed with liraglutide.

Biodegradable polymers are commonly used to design and to synthesize drug delivery systems (DDS). Among them, the most widely used are polyethylene glycol (PEG), poly(lactic acid) family and its copolymers with glycolic acid (Chen et al., 2003; Reddy, K. R., 2000). Conjugation of PEG to polymeric systems or drugs prevents recognition by various defense systems of organisms because of its potential for biomasking (Fontana et al., 2001; Huang et al., 2005; Perry et al., 2012). Furthermore they are biodegradable and biocompatible polymers which are approved by the US FDA for use in biomedical applications (Kamaly et al., 2016; Kapoor et al., 2015; Klose et al., 2008; Makadia and Siegel, 2011). In consequence some resent experimental researches in diabetes field are directed to the design of different types of DDS based on these co-polymers as a useful alternative for diabetes treatment (Cai et al., 2013; Guerreiro et al., 2012a, 2013). Although a long-acting release (LAR) system based on polymeric particle has been used for exenatide (Aroda and DeYoung, 2011; Liu et al., 2010; Pinelli and Hurren, 2011; Ryan et al., 2013b), to date there is no known sustained release formulation of either liraglutide or GLP-1 itself (Cai et al., 2013).

In this work, we explored the development of biocompatible polymeric microparticles loaded with either human GLP-1 (hGLP-1) and liraglutide and the comparative evaluation of *in vitro* kinetic release and *in vivo* pharmacologic evaluation of these products.

## 2 Material and Methods

### 2.1 Materials

Polyvinyl alcohol Mw 89 kDa-98kDa, >99% hydrolyzed (PVA; lote # MKBD2262V, Cat: 9002-89-5) was obtained from Sigma-Aldrich. Liraglutide (Victoza^®^, Lots # FS60K24 and FS6W992) was purchased at local drug store, and glucagon-like peptide-1 (GLP-1, 7-37, sequence “HAEGTFTSDVSSYLEGQAAKEFIAWLVKGRG”, > 95 % purit, lot #98664; Certificate of Analysis in *Supporting Material)* was purchased from Genemed Synthesis Inc, USA). Fluorescamine was obtained from Sigma-Aldrich (Cat #F9015). The branched PLGA-glucose (lots 0811001762 and Cat #5004-A) (Bodmer et al., 1992) and the linear PLGA 50:50 Lactide/Glicolide (lot 14007) were purchased from Purasorb. Linear PLGA 85:15 (Cat. #430471), 75:25 (Cat. #P1941) and 65:35 (Cat. #P2066) Lactide/Glicolide (lots #MK861113V, #089K134V and #050M1668V, respectively) were purchased from Sigma-Aldrich. PLA, PLA-PEG and PLA-PEG-F (fluorescein) were synthesized and characterized as described elsewhere (Icart et al., 2016). All other reagents were from analytical grade. All reactants were used as received.

### 2.2 Preparation of GLP-1 or Liraglutide-loaded polymeric microparticles by double-emulsion and solvent-evaporation procedure

Peptide loaded-polymeric microparticles were prepared using all the polymeric materials (PLA, PLA-PEG, PLA-PEG-F and PLGA) by using the double emulsion-solvent evaporation methodology (Allahyari and Mohit, 2015) with some modifications. In this methodology, we used 140 mg of the polymer solubilized in 1.00 mL of dichloromethane (DCM) (organic phase) and 1.2 mg or 3.6 mg of peptide solubilized in 200 μL of water (internal aqueous phase), targeting respectively a theoretical maximum load of 0.86 % and 2.6 % (peptide/polymer). First emulsion was formed mixing both immiscible phases under stirring at 20,000 rpm during 5 min, producing a w/o emulsion. Them, the resulting emulsion was transferred into a beaker containing 40 mL of PVA water solution (0.1 wt %) (external water phase). This system was kept under stirring at 20,000 rpm for 5 min, producing a w/o/w emulsion. Them the system was kept under mechanical stirring for 2 h at room temperature (25^°^C) to harden the microparticles by solvent evaporation. The peptide loaded-polymeric microparticles were then collected by centrifugation at 5,000 rpm and washed 3 times with water. Finally, the collected microparticles were freeze-drier by liofilization and stored at 12^°^C until use.

### 2.3 Characterization of polymeric microparticles

#### 2.3.1 Scanning electron microscopy

The morphologic analysis of the polymeric microparticles were carried out with a FEI-Quanta 259 Tungsten scanning electron microscope (at the analytical platform of CENABIO-UFRJ), using acceleration voltages of 12.5 kV. Samples were coated with gold and the materials were sampled by taking several images of various magnifications to ensure that the analysis was based on a representative region of the sample.

#### 2.3.2 Particle size distribution

The polymeric microparticles size distribution was accessed with the material dispersed in water and evaluated in a Mastersizer 2000 laser (Malvern instrument. Ltd. UK; at the analytical platform of EngePol, UFRJ) at room temperature.

#### 2.3.3 Peptide quantification

The quantification of the peptides was performed by derivatization with fluorescamine and fluorimetric analysis (Guerreiro et al., 2012b; Udenfriend et al., 1972). Briefly, an analytical curve (between 0.4 and 0.0065 μg/ μL) was performed by using the respective peptide (either liraglutide or hGLP-1) as standards. Samples were assessed in serial dilution in 96-well microplates (Corning A010993), in 50 mM sodium phosphate buffer pH 7.0 (final volume = 150 μL), added of 75 μL fluroescamine (500 μg/mL DMSO, for final concentration = 250 μg/mL) and read in a Spectramax M5 (Molecular Devices) after 5 min incubation at room temperature, using excitation at 390nm and emission at 480nm.

### 2.4 Encapsulation yield and encapsulation efficiency

Encapsulation yield was determined as the weight of the microparticles recovered (MPt) with respect to sum of all the starting material, peptide (PEPo) and polymer (POLo). The encapsulation yield of prepared microparticles was determined using Equation (1):

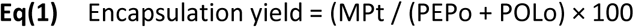

Encapsulation efficiency of peptides encapsulation was performed by quantification of the peptide by using the fluorescamine method (previously described) in the supernatant (PEPsn) after the microencapsulation process. This value was subtracted from the initial amount of peptides (PEPo) at the beginning of the process in order to calculate the amount of peptides incorporated to the microparticles. The percentage encapsulation efficiency of peptides was calculated with Equation 2:

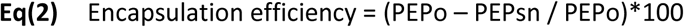

### 2.5 In *vitro* kinetic release of hGLP-1 or liraglutide-loaded microparticles

Peptide loaded-microparticles (containing 1 mg of peptide) were dispersed in 2 mL of phosphate buffer pH 7.2 and 100 mM of NaN_3_). Then 200 μl of the same microparticles suspension were placed in eppendorf tubes. The tubes were sealed and left under orbital agitation (150 rpm) at 37 ^°^C. At the indicated time interval (0, 1h, 24h, 48h, 4d, 8d, 15d, 20d, 25d, 30d and 45d) the tubes were removed from the shaker and they were centrifuged at 5,000 rpm at 4^0^C for 15 min, the supernatant was transferred to another eppendorf tube, flash frozen and kept at −20 ^o^C until use. Total peptide release was inferred by quantification by the fluorescamine method as described above.

### 2.6 Pharmacologic assays

The pharmacologic assays were approved by the Institutional Bioethics Committee on Animal Care and Experimentation at UFRJ (CCS, UFRJ, Protocol #057/17).

#### 2.6.1 Animals

Swiss male mice aged 7-8 weeks (25-28 g) were housed at a constant room temperature (22 ± 3^°^C) in a 12 hours light/dark cycle and fed *ad libitum* with standard diet purchased from Nuvilab (DSPS803 Cat # 92-33634).

#### 2.6.2 Single dose of hGLP-1 or liraglutide-loaded polymeric microparticles

The mice were randomly distributed into three groups (n=5/groups) and left 7days of housing and adaptation before subjected to experimentation. The experimental design took into account the idealization of a hypothetical linear, constant flux of peptide release from the particles resulting in an approximate dose of 200 μg peptide/body weight/day, during 40 days (based on the estimate extension of the kinetic *in vitro* release). Based on this estimation we achieved a dose of about 5 μg peptide/animal (of approximately 25 g body weight).

The suspension (in 0.9 % NaCl saline) contained 40 mg microparticles/100 μL, either dummy (formulated without peptide) or loaded with peptide (either liraglutide or hGLP-1, at 5.6 μg peptide / mg particles), corresponding to a dose of about 224 μg peptide / 25 g body weight (or 8.96 mg peptide / kg body weight). Each animal received 100 μL suspension/25 g body weight. The formulation was administer by subcutaneous injection (over the shoulders, into the loose skin over the neck) using a standard 29 gauge needle (BD™).

#### 2.6.3 Measurement of blood glucose and weight

Mice were weighing regularly and had they glycemia measured weekly (at about 2 pm); with a portable glucometer ACCU-CHEK, Cat # TYP: 05680456003), both in baseline and after microparticle administration.

#### 2.6.4 Statistical analysis

Data are the Means ± Standard Media Error (SME). Weight change, glucose change, food and water intake and feces and urine production were compared across groups using two-way ANOVA, with Bonferroni multiple comparison subtests. Graphs and statistical anaylsis were obtained using GraphPadPrism 5.0 (GraphPad Software Inc.).

## 3 Results

### 3.1 Preparation of hGLP-1- or liraglutide-loaded polymeric microparticles

The preparation of polymeric microparticles by emulsion and solvent evaporation is a satisfactory procedure for the entrapment of small compounds aimed their chemical of physical stabilization and sustained release (“Emulsion Solvent Evaporation Microencapsulation Review | Emulsion | Polyethylene Glycol,” n.d.; R and V, 2015; Yüksel and Baykara, 1997). A large set of variables can be explored in their preparation according to the compound characteristics and aiming to achieve the desired morphology, size, polidispersity, release profile, yield and entrapment efficiency (Tu and Lee, 2012). We choose using the double emulsion and solvent evaporation approach in the preparation of hGLP-1 and liraglutide loaded microparticles since this procedure is one of the most useful methods for entrapping water-soluble substance, such as proteins and peptide. The high hydrophilicity of these molecules favors their quantitative introduction into the internal aqueous phase which results in an increased encapsulation efficiency in comparison with some others emulsifications procedures such as single-emulsion and solvent evaporation. In addition, the use of proteins induces a stabilizing effect of the emulsions formed, which contributes to the success of the double emulsion process and loading.

We used a set of 8 polymers in this study, covering the classes of PLA, PLA-PEG and PLGA, which are the most commonly used biocompatible polymers for therapeutic purpose (Han et al., 2016; Kamaly et al., 2016; Makadia and Siegel, 2011). The characteristics of the polymers used here and the resulting microparticles are detailed in Table 1. The double emulsion and solvent evaporation process prepared with a ratio of 0.86 % peptide/polymer yielded microparticles with satisfactory narrow polydispersity of size distribution (Pd = 0.2-0.8). The microparticles had a mean diameter of 44.1± 10.8 and 47.8 ±14.7 μm for hGLP-1 and liraglutide respectively, well below the safety cutoff of 100 μm (due to overall risks of embolization) (Lee and Henthorn, 2012). The microparticles was in the range of 57-76% for hGLP-1 and 82-95% for liraglutide. Peptide encapsulation efficiency ranged from 47-68% to hGLP-1 and 54-74% for liraglutide.

**Table 1:** **Characteristic of the polymer and resulting microparticles formulated with hGLP1 or liraglutide (0.86 % load, peptide/polymer).**

| Materials |  |  | hGLP1-loaded microparticles |  |  |  | Liraglutide-loaded microparticles |  |  |  |
| --- | --- | --- | --- | --- | --- | --- | --- | --- | --- | --- |
| Polymer | L/G* | Mw (kda) | Size (µm) | Pd | Yield (%) | E.E** (%) | Size (µm) | Pd | Yield (%) | E.E (%) |
| PLGA (A) | 50:50 | 60-80 | 48.2 | 0.3 | 75.1 | 52.0 | 53.0 | 0.3 | 95.5 | 62.2 |
| PLGA (B) | 85:15 | 50-75 | 56.2 | 0.4 | 60.5 | 54.5 | 57.2 | 0.4 | 82.3 | 74.6 |
| PLGA (C) | 75:25 | 66-77 | 61.1 | 0.4 | 57.9 | 59.7 | 68.5 | 0.2 | 84.2 | 61.9 |
| PLGA (D) | 65:35 | 40-75 | 43.3 | 0.3 | 64.8 | 56.9 | 40.9 | 0.3 | 86.6 | 64.2 |
| PLGA (E) | 55:45 | 50-90 | 48.4 | 0.3 | 73.5 | 57.3 | 66.9 | 0.4 | 82.7 | 54.2 |
| PLA | n/a | 10-20 | 30.7 | 0.6 | 76.8 | 53.9 | 32.3 | 0.4 | 91.2 | 61.4 |
| PLA-PEG | n/a | 10-20 | 28.1 | 0.3 | 59.6 | 47.5 | 27.1 | 0.4 | 87.7 | 66.5 |
| PLA-PEG-F | n/a | 10-20 | 37.5 | 0.8 | 75.3 | 43.2 | 37.2 | 0.6 | 92.8 | 73.8 |

\*L/G: Lactide/glicolide composition.
\*\*Encapsulation efficiency.

The particle size of the microparticles obtained in this work were mainly influence by the average molecular weight of the polymer. Microparticles which were prepared with PLA or PLA-PEG showed an smaller particle sized than microparticles prepared with the co-polymers of PLGA, (**Table 1**). The morphological characterization of prepared microparticles containing hGLP-1 (**Fig. 1**) and liraglutide (**Fig. 2**) by SEM revealed the influence of the polymer composition on the morphological surface of microparticles. Microparticles which were prepared using PLA, showed a smoother surface with a few randomly distributed small pores. In turn, microparticles prepared with co-polymer of PLA-PEG showed a rough surface with a few randomly distributed small pores. This might be attributed to the known effect of PEG in the induction of small pores in the surfaces of polymeric microparticles. During the precipitation step, the PEG branches of the diblock copolymer are known to be orientated toward the internal and surrounding aqueous phase forming a sponge-like structure. This phenomenon was also reported for microparticles based on blends of PLGA/PLA and PEG and also applies to microparticles composed of diblock polymer (Essa et al., 2010; Lochmann et al., 2010). The amount of glicolide into the composition of the PLGA was also likely to influence the morphological characteristics of the prepared microparticles.

**Figure 1.**
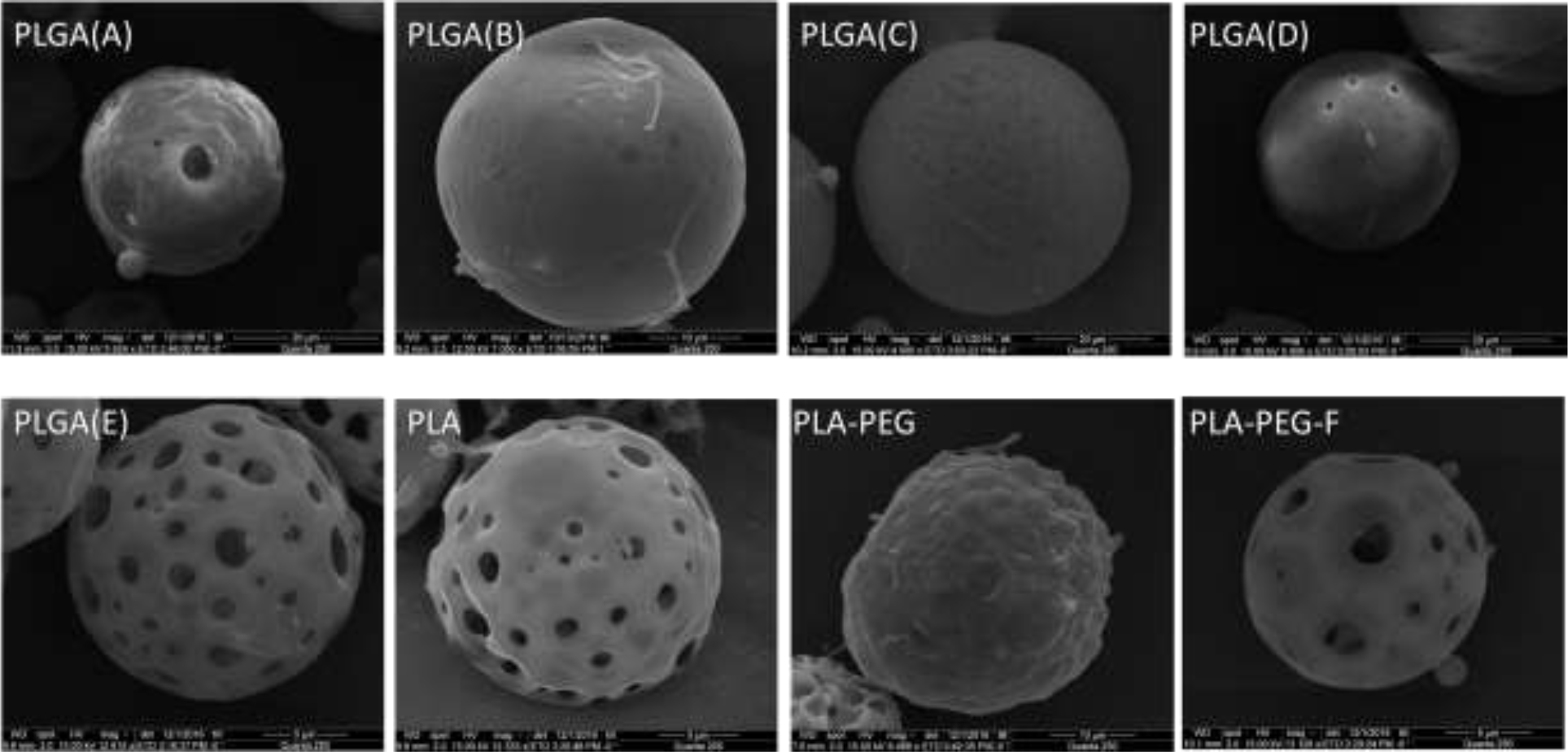
Morphologic characterization of the h-GLP-1 loaded microparticles by scanning electron microscopy (SEM). The hGLP-1-loaded microparticles were formulated used the different PLGA commercial co-polymers (A, B, C, D and D) and the synthetic polymers (PLA, PLA-PEG and PLA-PEG-F) as depicted in the respective panels. The detailed information about the methods and the polymers cam be found in the Material and Methods section and in **Table 1.**

**Figure 2.**
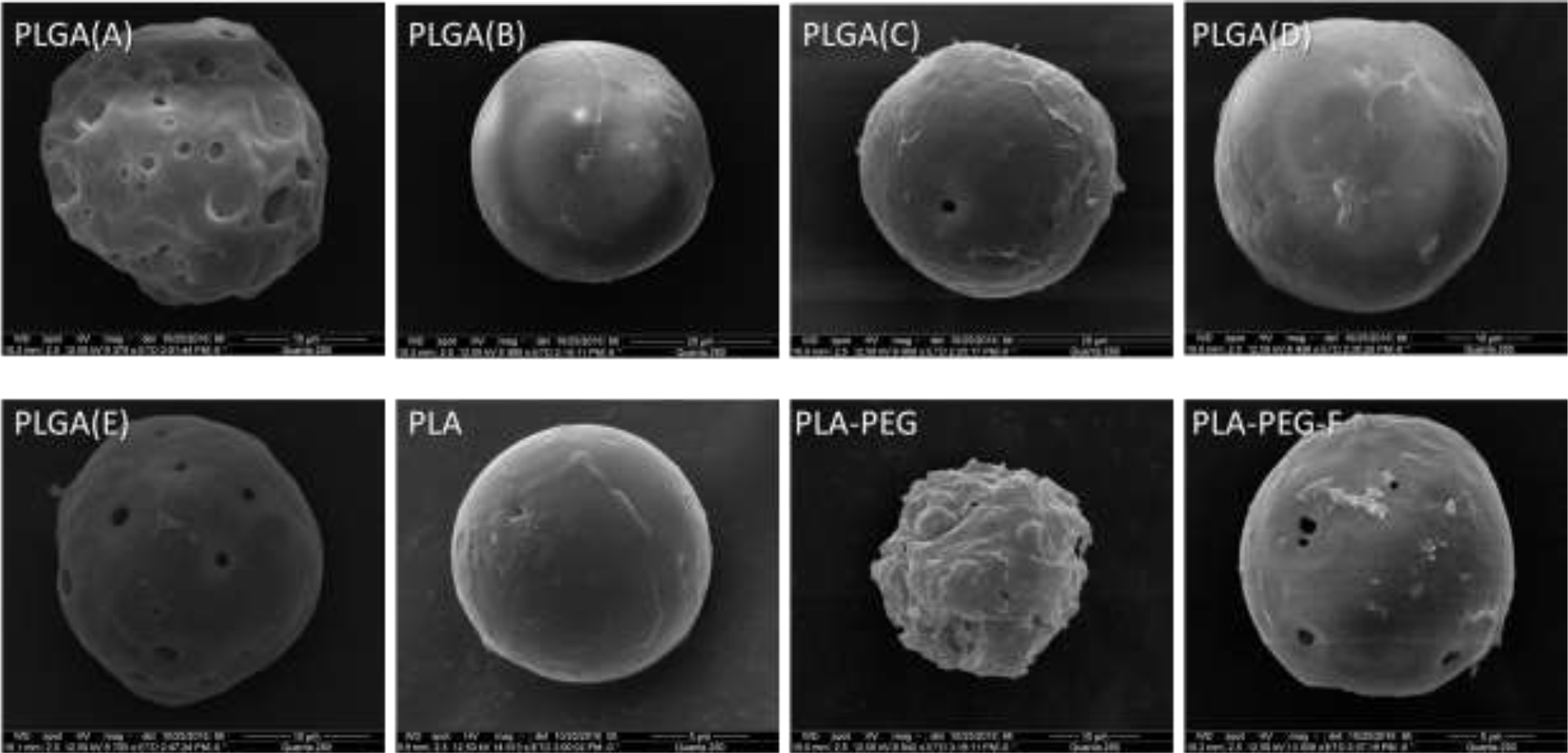
Morphologic characterization of the liraglutide loaded microparticles by scanning electron microscopy (SEM). The liraglutide-loaded microparticles were formulated used the different PLGA commercial co-polymers (A, B, C, D and D) and the synthetic polymers (PLA, PLA-PEG and PLA-PEG-F) as depicted in the respective panels. The detailed information about the methods and the polymers cam be found in the Material and Methods section and in **Table 1**.

Microparticles prepared using PLGA-A and PLGA-E with high glicolide composition higher that those based on PLGA-B, PLGA-C and PLGA-D showed a more irregular surface and several apparent large pores. Instead, the microparticles based on PLGA with lower glicolide content were more regular and showed smoother surface with fewer pores.

Microparticles prepared with a 3 times higher load (2.6 % liraglutide/polymer) resulted in microparticles similar in morphology (Fig. S1) and particle size, encapsulation efficiency and encapsulation (Table S1) to the microparticles preparations formulated with 0.86 % load (Table 1; Fig. 2). These results are in agreement with those reported for different types of peptides microencapsulation by double emulsion and solvent evaporation (Parikh et al., 2003; Ravi et al., 2016; Xu et al., 2006; Yang and Owusu-Ababio, 2000).

### 3.2 *in vitro* kinetic release profile of hGLP-1 and liraglutide from microparticles

We next evaluated the *in vitro* kinetic release profile of hGLP-1 and liraglutide peptides from the polymeric microparticles. The **Figure 3** shows the results of the kinetic release assay performed with hGLP-1 (**Fig. 3a** and **Fig. 3b**) and liraglutide (**Fig. 3c** and **Fig. 3d**) for 45 days in phosphate buffer pH 7.4 at 37^0^C. Within the first days an initial ‘burst’ of peptide release took place for hGLP-1 (~30% of total peptide) and liraglutide (~10% of total peptide). A hyperbolic profile comprised by a burst phase followed by a slower kinetic release between approximately day 5 up to the end of the study is observed for the hGLP-1-loaded microparticles, with either PLGA (**Fig. 3c**) or PLA (**Fig. 3d**), Instead, the *in vitro* kinetic release profile liraglutide from the microparticles showed a sigmoidal pattern after the small burst phase, with a plateau up to about day 20, followed by an exponential phase for the rest of the assay, for both PLGA (**Fig. 3a**) and PLA (**Fig. 3b**). This dissimilar kinetic release behavior between microparticles loaded with hGLP-1 or the palmitoylated analogue liraglutide suggests a deterministic effect of the fatty acid moiety of the conjugate on the resulting microparticles.

**Figure 3.**
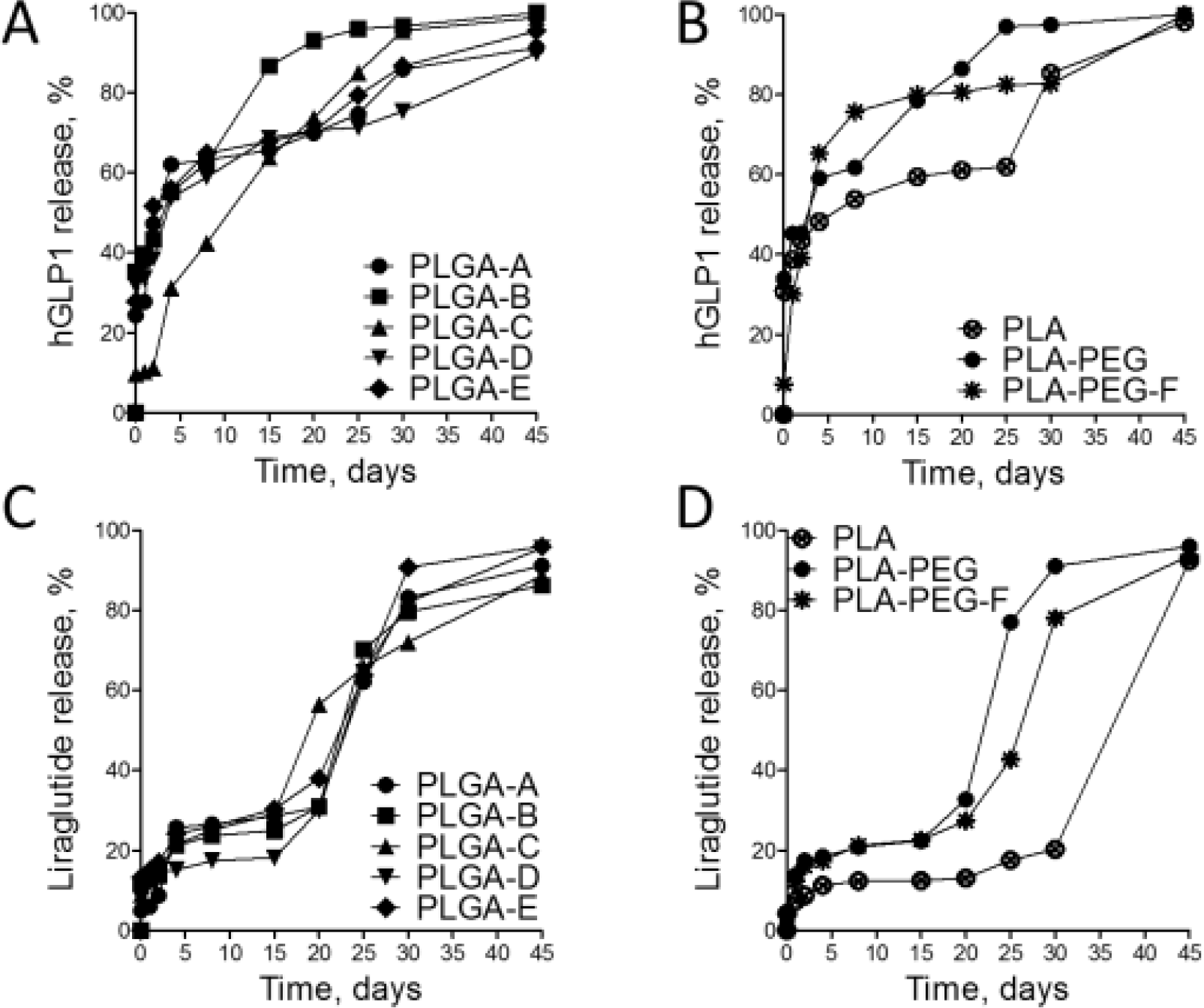
*In vitro* kinetic release profile of hGLP-1 and liraglutide from peptide-loaded microparticles. The microparticles loaded with hGLP-1 (**A** and **B**) or liraglutide (**C** and **D**) were prepared using the PLGA (**A** and **C**) and PLA (**B** and **D**) depicted in **Table 1.** The release was performed in phosphate solution pH 7.4, 37°C, under orbital agitation set at 75 rpm. The peptide (hGLP-1 or liraglutide) released from the polymeric microparticles at each time interval was quantified from supernatant harvested after centrifugation (4°C, 15 min, 5,000rpm) of aliquots and them quantified by the fluorescamine procedure. Details in the *Material and Methods* section.

### 3.3 Pharmacologic evaluation of hGLP-1- and liraglutide-loaded microparticles

We have further evaluated the *in vivo* effect of these formulations of hGLP-1 and liraglutide polymeric microparticles. The microparticles produced with PLA resulted in a faster kinetic release profile compared to the extended release obtained with the microparticles formulated with the PLGA polymers. The polymers PLGA-A and PLGA-E have both the more equivalente amount of lactide:glicolide, while PLGA-B and PLGA-C were higher in lactide – such as the PLA only. However, the particles obtained with PLGA-A and PLGA-E were irregular in morphology and were hollow. Instead, the PLGA-D showed uniformity in particle surface when used in preparation of microparticles with both peptides. Based on these features, we choose the PLGA-D as a polymer for the pharmacologic assay of the peptide-loaded microparticles (hGLP-1 and liraglutide), along with control with polymer-only microparticles.

Swiss male mice aged 7-8 weeks (25-28 g) were randomly distributed in three groups (n=5/group) and they received each subcutaneously a single dose of unloaded, hGLP-1-loaded or liraglutide-loaded polymeric microparticles. The group receiving microparticles containing either hGLP-1 or liraglutide showed a decline in the rate of body weight gain after at 9 days period with no significative effect being observed. From day 10 up to day 25 after dosing, a significative difference was observed, both for body weight (Fig. 4a) and reduction in glycemia (Fig. 4b). Collectively, these results demonstrate the similar long-term effectiveness of both unmodified hGLP-1 and the analogue liraglutide in the present *in vivo* pharmacological assays.

**Figure 4:**
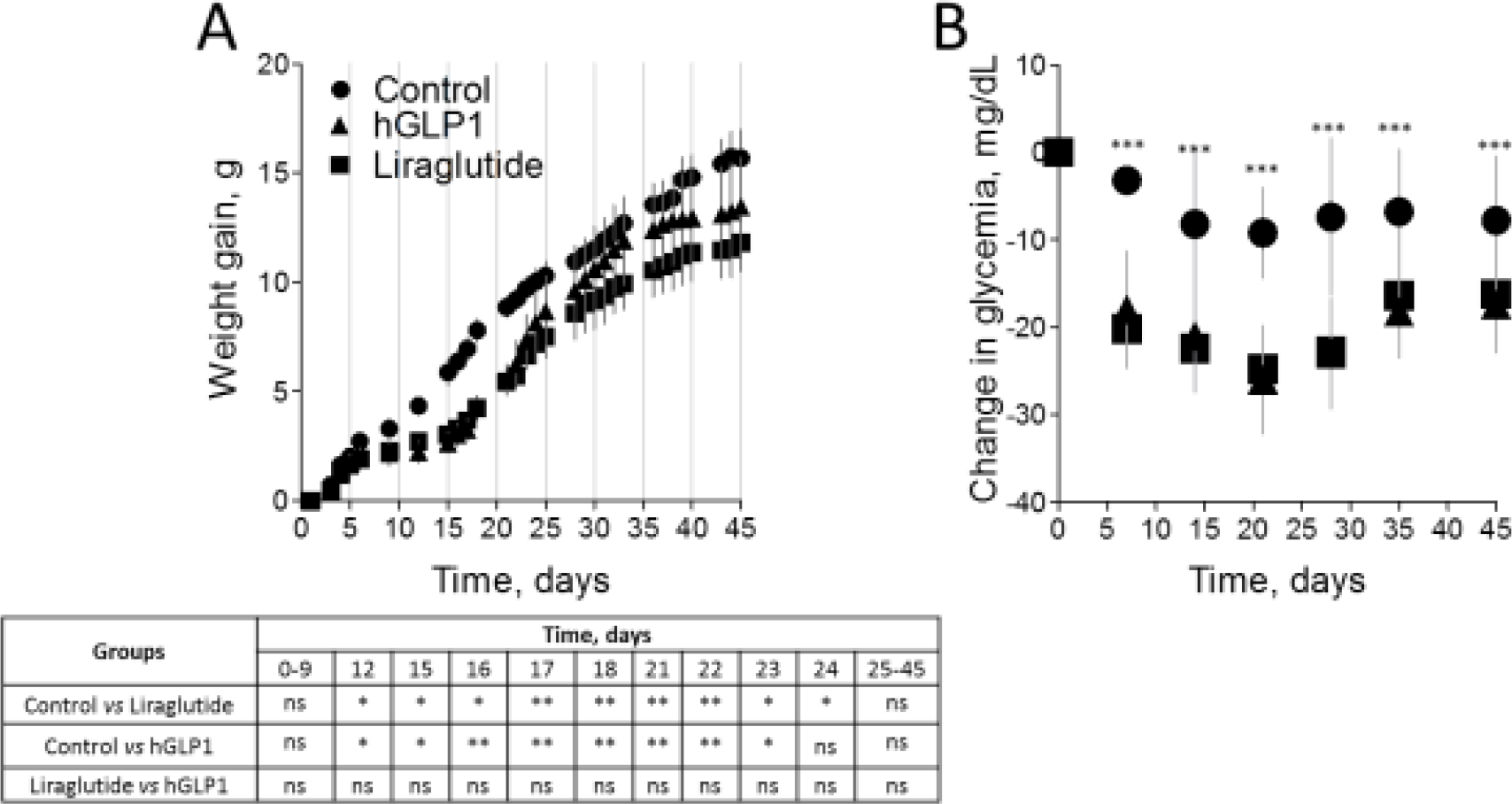
Pharmacologic evaluation of hGLP-1-loaded or liraglutide-loaded microparticles. Swiss male mice aged 7-8 weeks (n=5/groups) received a single dose of microparticles formulation (either with no peptide, loaded with hGLP-1 or liraglutide) and were monitored for 45 days for change in body weight (A) and glycemia (B). ^*^p < 0.05, ^**^p < 0.01, and ^***^p < 0.001.

## 4 Discussion

In this work we have shown that the well-established pharmaceutical system of long-acting release based on microparticles developed with biocompatible polymers could be used for the formulation of the regular human GLP-1 and its analogue liraglutide. The effect of reducing body weight and glycemia in mice for over 3 weeks provided evidence for the *in vivo* effect of these formulations, which upon further optimization could be considered as an alternative approach in the therapeutic portfolio for diabetes, obesity and comorbidities.

The very short serum half-life of hGLP-1 of about 2 min has been hampering its direct therapeutic use, and motivated the development of other GLP-1 receptor agonists such as its analogue liraglutide. However, upon formulation into polymeric microparticles with same PLGA polymer, these products resulted in close *in vivo* effect of both hGLP-1 and liraglutide (Fig. 4), suggesting similar potency, overall release and absorption flux, although further other pharmacokinetic, pharmacodynamics and toxicological investigation would be desirable to establish a better foundations for the understanding of those systems.

From our kinetic release *in vitro* we could observe a multi-pattern profile, mainly in three phases:

i. an initial fast release (defined as “burst release”), starting within the first hours and can last for a few days, and is most commonly attributed to drug adhered to the external wall of the microparticles,
ii. an slow release phase, which depends mainly on the degradation kinetics of the polymeric matrix used to prepare the microparticles, and
iii. the last accelerated release, which depends on the microsphere diameter and drug diffusion or solubility (Kamaly et al., 2016).

In this study we observed a deterministic effect of both the polymer (PLGA versus PLA) and the nature of the entrapped compound (hGLP-1 *versus* its palmitoylated analogue liraglutide) in the peptide release profile from the microparticles. Hence microparticles based on PLGA showed a faster hGLP-1 and liraglutide release compared to PLA, most likely due to the increase in the ductility of the PLGA material (Song et al., 2011). In the case of the PLGA microparticles, no large effect of the proportion of lactide and glicolide was observed on the peptides release profiles.

*In vitro* polymer degradation is generally considered as heterogeneous process. First degradation probably occurs on the surface due to the polymer water intake, however the higher concentration of carbonyl groups in the center of the particle helps the self-catalyzing degradation. This behavior is common during the degradation of aliphatic polyesters (Ruan and Feng, 2003). Furthermore it was observed a faster release of hGLP-1 compared to liraglutide, which may be attributed to the enhanced aqueous solutiliby of hGLP-1 compared to the fatty acid acylated liraglutide. Finally, liraglutide loaded-microparticles prepared using the triple load of peptide (2.5 % instead of 0.86 %) showed both similar kinetic release (**Supporting Material**).

The relationships between GLP-1 or GLP-1-analogous with appetite and weight maintenance was previously reported by several authors (Aroda and DeYoung, 2011; DeFronzo et al., 2005, p. 1; Larsen et al., 2001; Madsbad, 2009; Madsbad et al., 2004). Some evidence demonstrates that GLP-1 reduces body weight when administered by intracerebroventricular route in animals (Ronveaux et al., 2015; Tang-Christensen et al., 1996; Turton et al., 1996) and by subcutaneous route in humans (Abbott et al., 2005; Rüttimann et al., 2009; Talsania et al., 2005; Williams et al., 2009). It´s known that they have action on the gastrointestinal tracts as well as the direct regulation of appetite which provoke inhibition of gastric secretion and motility. Some recent research suggests that long-acting analogs of the GLP-1 could be a useful alternative to obesity treatment (Cai et al., 2013). In consequence the GLP-1 receptor agonist was recently approved by the US Food and Drug Administration as an obesity treatment option, showing interesting results in human reducing and sustaining body weight loss (“FDA Approves Liraglutide (Saxenda) for Weight Loss,” n.d.; Isaacs et al., 2016). In addition, Figure 3b also confirms the effects of the released GLP-1 or Liraglutide from microparticles on the blood glucose levels of the treated groups. In which an important decline of blood glucose levels was observed with statistical significance difference as compared with control group (p value = 0.0003). No statistical significance difference was observed between both treated groups. This effect was sustained during at least fourth week of study. From this moment and to the end of the experiment a tendency to increase of this parameter was observed. Currently several pre-clinical and clinical studies demonstrated the role of GLP-1 on the blood glucose homeostasis due to its capacity to slow gastric emptying, to enhance pancreatic insulin secretion, and to suppress pancreatic glucagon secretion (Farilla et al., 2003; Gough, 2012, p. 1; Madsbad, 2009, p. 1; Madsbad et al., 2004, p. 1; Minze et al., 2013, p. 1; Ronveaux et al., 2015).

In summary, the present study demonstrated the effective ability of trapping hGLP-1 and its analogue liraglutide into biocompatible polymeric microparticles, resulting in an extended release profile and overcoming the limitations of the very short half-life of hGLP-1.

## 5 Conclusions

The emulsion and solvent evaporation procedure allowed obtaining hGLP-1 or liraglutide loaded-microparticles with satisfactory peptide encapsulation yield and microparticles recovery. The peptide-loaded microparticles showed an extended kinetic release profile spanning over 5 weeks *in vitro* with pharmacological effect sustained for over 3 weeks, even for the hGLP-1 known for its short half-life. These results pave a possibility for new long-acting release formulation based either on the unmodified hGLP-1 or the well-established analogue liraglutide.

## Acknowledgments

We would like to thank pharmacysts Thayna Sisnande, Dayana Cabral and Celimar Sinesia for helpful assistance with *in vivo* assays, to Dr. Sergio Luiz Cordeiro (CENABIO-UFRJ), Prof. Marcos Lopes Dias (IMA, UFRJ), and Dr. Larissa Leite de Almeida Carvalho (EngePol, COPPE, UFRJ) for excellent technical support, and to Laboratory Hertha-Meyer (IBCCF, UFRJ), CENABIO (UFRJ), EngePol (UFRJ) and IMA (UFRJ) for providing access to their analytical facilities. This work was supported by the Coordenação de Aperfeiçoamento de Pessoal de Niveĺ Superior (CAPES), Conselho Nacional de Desenvolvimento Científico e Tecnológico (CNPq), Fundação de Amparo à Pesquisa do Estado do Rio de Janeiro Carlos Chagas Filho (FAPERJ). The funding agencies had no role in the study design, data collection and analysis, or decision to publish or prepare of the manuscript.

## Conflict of interest

The authors have no financial conflicts of interest with the contents of this article. LMTRL is a participant in patent applications by the UFRJ on controlled release of peptides unrelated to the present work.

## Author Contributions

Conceived and designed the experiments: LPI, FGSJ, LMTRL

Performed the experiments: LPI, FGSJ, LMTRL

Analyzed the data: LPI, FGSJ, LMTRL

Contributed reagents/materials/analysis tools: LPI, FGSJ, LMTRL

Wrote the manuscript: LPI, FGSJ, LMTRL

## References

Abbott, C.R., Monteiro, M., Small, C.J., Sajedi, A., Smith, K.L., Parkinson, J.R.C., Ghatei, M.A., Bloom, S.R., 2005. The inhibitory effects of peripheral administration of peptide YY(3-36) and glucagon-like peptide-1 on food intake are attenuated by ablation of the vagal-brainstem-hypothalamic pathway. Brain Res. 1044, 127–131. https://doi.org/10.1016/j.brainres.2005.03.011

Agersø, H., Vicini, P., 2003. Pharmacodynamics of NN2211, a novel long acting GLP-1 derivative. Eur. J. Pharm. Sci. Off. J. Eur. Fed. Pharm. Sci. 19, 141–150.

Ahrén, B., Schmitz, O., 2004. GLP-1 receptor agonists and DPP-4 inhibitors in the treatment of type 2 diabetes. Horm. Metab. Res. Horm. Stoffwechselforschung Horm. Metab. 36, 867–876. https://doi.org/10.1055/s-2004-826178

Allahyari, M., Mohit, E., 2015. Peptide/protein vaccine delivery system based on PLGA particles. Hum. Vaccines Immunother. 12, 806–828. https://doi.org/10.1080/21645515.2015.1102804

Aroda, V.R., DeYoung, M.B., 2011. Clinical implications of exenatide as a twice-daily or once-weekly therapy for type 2 diabetes. Postgrad. Med. 123, 228–238. https://doi.org/10.3810/pgm.2011.09.2479

Bethel, M.A., Patel, R.A., Merrill, P., Lokhnygina, Y., Buse, J.B., Mentz, R.J., Pagidipati, N.J., Chan, J.C., Gustavson, S.M., Iqbal, N., Maggioni, A.P., Öhman, P., Poulter, N.R., Ramachandran, A., Zinman, B., Hernandez, A.F., Holman, R.R., 2018. Cardiovascular outcomes with glucagon-like peptide-1 receptor agonists in patients with type 2 diabetes: a meta-analysis. Lancet Diabetes Endocrinol. 6, 105–113. https://doi.org/10.1016/S2213-8587(17)30412-6

Bodmer, D., Kissel, T., Traechslin, E., 1992. Factors influencing the release of peptides and proteins from biodegradable parenteral depot systems. J. Controlled Release 21, 129–137. https://doi.org/10.1016/0168-3659(92)90014-I

Brunton, S., 2014. GLP-1 receptor agonists vs. DPP-4 inhibitors for type 2 diabetes: is one approach more successful or preferable than the other? Int. J. Clin. Pract. 68, 557–567. https://doi.org/10.1111/ijcp.12361

Cai, Y., Wei, L., Ma, L., Huang, X., Tao, A., Liu, Z., Yuan, W., 2013. Long-acting preparations of exenatide. Drug Des. Devel. Ther. 7, 963–970. https://doi.org/10.2147/DDDT.S46970

Chen, C.C., Chueh, J.Y., Tseng, H., Huang, H.M., Lee, S.Y., 2003. Preparation and characterization of biodegradable PLA polymeric blends. Biomaterials 24, 1167–1173.

DeFronzo, R.A., Ratner, R.E., Han, J., Kim, D.D., Fineman, M.S., Baron, A.D., 2005. Effects of Exenatide (Exendin-4) on Glycemic Control and Weight Over 30 Weeks in Metformin-Treated Patients With Type 2 Diabetes. Diabetes Care 28, 1092–1100. https://doi.org/10.2337/diacare.28.5.1092

Emulsion Solvent Evaporation Microencapsulation Review | Emulsion | Polyethylene Glycol [WWW Document], n.d.. Scribd. URL https://es.scribd.com/document/284394619/Emulsion-Solvent-Evaporation-Microencapsulation-Review (accessed 6.23.17).

Essa, S., Rabanel, J.M., Hildgen, P., 2010. Effect of polyethylene glycol (PEG) chain organization on the physicochemical properties of poly(d, l-lactide) (PLA) based nanoparticles. Eur. J. Pharm. Biopharm. 75, 96–106. https://doi.org/10.1016j.ejpb.2010.03.002

Farilla, L., Bulotta, A., Hirshberg, B., Li Calzi, S., Khoury, N., Noushmehr, H., Bertolotto, C., Di Mario, U., Harlan, D.M., Perfetti, R., 2003. Glucagon-Like Peptide 1 Inhibits Cell Apoptosis and Improves Glucose Responsiveness of Freshly Isolated Human Islets. Endocrinology 144, 5149–5158. https://doi.org/10.1210/en.2003-0323

FDA Approves Liraglutide (Saxenda) for Weight Loss [WWW Document], n.d. . Medscape. URL http://www.medscape.com/viewarticle/837147 (accessed 10.24.17).

Fontana, G., Licciardi, M., Mansueto, S., Schillaci, D., Giammona, G., 2001. Amoxicillin-loaded polyethylcyanoacrylate nanoparticles: influence of PEG coating on the particle size, drug release rate and phagocytic uptake. Biomaterials 22, 2857–2865.

Gough, S.C.L., 2012. Liraglutide: from clinical trials to clinical practice. Diabetes Obes. Metab. 14 Suppl 2, 33–40. https://doi.org/10.1111/j.1463-1326.2012.01576.x

Guerreiro, L.H., Da Silva, D., Ricci-Junior, E., Girard-Dias, W., Mascarenhas, C.M., Sola-Penna, M., Miranda, K., Lima, L.M.T.R.,2012a. Polymeric particles for the controlled release of human amylin. Colloids Surf. B Biointerfaces 94, 101–106.https://doi.org/10.1016/j.colsurfb.2012.01.021

Guerreiro, L.H., Girad-Dias, W., Miranda, K.R. de, Lima, L.M.T.R.,2012b. A fluorescence-based assay for octreotide in kinetic release from depot formulations. Quím. Nova 35, 1025–1029. https://doi.org/10.1590/S0100-40422012000500029

Guerreiro, L.H., Guterres, M.F.A.N., Melo-Ferreira, B., Erthal, L.C.S., da Silva Rosa, M., Lourenço, D., Tinoco, P., Lima, L.M.T.R., 2013. Preparation and Characterization of PEGylated Amylin. AAPS PharmSciTech 14, 1083–1097. https://doi.org/10.1208/s12249-013-9987-4

Han, F.Y., Thurecht, K.J., Whittaker, A.K., Smith, M.T., 2016. Bioerodable PLGA-Based Microparticles for Producing Sustained-Release Drug Formulations and Strategies for Improving Drug Loading. Front. Pharmacol. 7. https://doi.org/10.3389/fphar.2016.00185

Holst, J.J., 2007. The physiology of glucagon-like peptide 1. Physiol. Rev. 87, 1409–1439. https://doi.org/10.1152/physrev.00034.2006

Huang, M., Wu, W., Qian, J., Wan, D.-J., Wei, X.-L., Zhu, J.-H., 2005. Body distribution and in situ evading of phagocytic uptake by macrophages of long-circulating poly (ethylene glycol) cyanoacrylate-co-n-hexadecyl cyanoacrylate nanoparticles. Acta Pharmacol.Sin. 26, 1512–1518. https://doi.org/10.1111/j.1745-7254.2005.00216.x

Icart, L.P., Fernandes, E., Agüero, L., Ramón, J., Zaldivar, D., Dias, M.L., 2016. Fluorescent microspheres of poly(ethylene glycol)-poly(lactic acid)-fluorescein copolymers synthesized by Ugi four-component condensation. J. Appl. Polym. Sci. 133, n/a-n/a. https://doi.org/10.1002/app.42994

Isaacs, D., Prasad-Reddy, L., Srivastava, S.B., 2016. Role of glucagon-like peptide 1 receptor agonists in management of obesity. Am. J. Health-Syst. Pharm. AJHP Off. J. Am. Soc. Health-Syst. Pharm. 73, 1493–1507. https://doi.org/10.2146/ajhp150990

Kamaly, N., Yameen, B., Wu, J., Farokhzad, O.C., 2016. Degradable Controlled-Release Polymers and Polymeric Nanoparticles: Mechanisms of Controlling Drug Release. Chem. Rev. 116, 2602–2663. https://doi.org/10.1021/acs.chemrev.5b00346

Kapoor, D.N., Bhatia, A., Kaur, R., Sharma, R., Kaur, G., Dhawan, S., 2015. PLGA: a unique polymer for drug delivery. Ther. Deliv. 6, 41–58. https://doi.org/10.4155/tde.14.91

Kim, H.J., Kim, D.J., 2017. Lessons from a cardiovascular outcome trial with liraglutide in type 2 diabetes. J. Diabetes Investig. 8, 431–433. https://doi.org/10.1111/jdi.12607

Klose, D., Siepmann, F., Elkharraz, K., Siepmann, J., 2008. PLGA-based drug delivery systems: Importance of the type of drug and device geometry. Int. J. Pharm., Special Issue in Honor of Prof. Tsuneji Nagai 354, 95–103. https://doi.org/10.1016/j.ijpharm.2007.10.030

Larsen, P.J., Fledelius, C., Knudsen, L.B., Tang-Christensen, M., 2001. Systemic Administration of the Long-Acting GLP-1 Derivative NN2211 Induces Lasting and Reversible Weight Loss in Both Nogrmal and Obese Rats. Diabetes 50, 2530–2539. https://doi.org/10.2337/diabetes.50.11.2530

Lee, S., Henthorn, D., 2012. Materials in Biology and Medicine. CRC Press.

Liu, B., Dong, Q., Wang, M., Shi, L., Wu, Y., Yu, X., Shi, Y., Shan, Y., Jiang, C., Zhang, X., Gu, T., Chen, Y., Kong, W., 2010. Preparation, characterization, and pharmacodynamics of exenatide-loaded poly(DL-lactic-co-glycolic acid) microspheres. Chem. Pharm. Bull. (Tokyo) 58, 1474–1479.

Lochmann, A., Nitzsche, H., von Einem, S., Schwarz, E., Mäder, K., 2010. The influence of covalently linked and free polyethylene glycol on the structural and release properties of rhBMP-2 loaded microspheres. J. Controlled Release 147, 92–100. https://doi.org/10.1016/j.jconrel.2010.06.021

Madsbad, S., 2009. Exenatide and liraglutide: different approaches to develop GLP-1 receptor agonists (incretin mimetics) - preclinical and clinical results. Best Pract. Res. Clin. Endocrinol. Metab., Incretins 23, 463–477. https://doi.org/10.1016/j.beem.2009.03.008

Madsbad, S., Schmitz, O., Ranstam, J., Jakobsen, G., Matthews, D.R., 2004. Improved Glycemic Control With No Weight Increase in Patients With Type 2 Diabetes After Once-Daily Treatment With the Long-Acting Glucagon-Like Peptide 1 Analog Liraglutide (NN2211) A 12-week, double-blind, randomized, controlled trial. Diabetes Care 27, 1335–1342. https://doi.org/10.2337/diacare.27.6.1335

Makadia, H.K., Siegel, S.J., 2011. Poly Lactic-co-Glycolic Acid (PLGA) as Biodegradable Controlled Drug Delivery Carrier. Polymers 3, 1377–1397. https://doi.org/10.3390/polym3031377

Marso, S.P., Daniels, G.H., Brown-Frandsen, K., Kristensen, P., Mann, J.F.E., Nauck, M.A., Nissen, S.E., Pocock, S., Poulter, N.R., Ravn, L.S., Steinberg, W.M., Stockner, M., Zinman, B., Bergenstal, R.M., Buse, J.B., 2016. Liraglutide and Cardiovascular Outcomes in Type 2 Diabetes. N. Engl. J. Med. 375, 311–322. https://doi.org/10.1056/NEJMoa1603827

Minze, M.G., Klein, M.S., Jernigan, M.J., Wise, S.L., Frugé, K., 2013. Once-weekly exenatide: an extended-duration glucagon-like peptide agonist for the treatment of type 2 diabetes mellitus. Pharmacotherapy 33, 627–638. https://doi.org/10.1002/phar.1240

Parikh, R.H., Parikh, J.R., Dubey, R.R., Soni, H.N., Kapadia, K.N., 2003. Poly(D,L-Lactide-Co-Glycolide) microspheres containing 5-fluorouracil: Optimization of process parameters. AAPS PharmSciTech 4, 14–21. https://doi.org/10.1208/pt040213

Perry, J.L., Reuter, K.G., Kai, M.P., Herlihy, K.P., Jones, S.W., Luft, J.C., Napier, M., Bear, J.E., DeSimone, J.M., 2012. PEGylated PRINT Nanoparticles: The Impact of PEG Density on Protein Binding, Macrophage Association, Biodistribution, and Pharmacokinetics. Nano Lett. 12, 5304–5310. https://doi.org/10.1021/nl302638g

Pinelli, N.R., Hurren, K.M., 2011. Efficacy and safety of long-acting glucagon-like peptide-1 receptor agonists compared with exenatide twice daily and sitagliptin in type 2 diabetes mellitus: a systematic review and meta-analysis. Ann. Pharmacother. 45, 850–860. https://doi.org/10.1345/aph.1Q024

Ponzani, P., 2013. Long-term effectiveness and safety of liraglutide in clinical practice. Minerva Endocrinol. 38, 103–112.

R, K.N., V, S., 2015. Microsphere: A Brief Review. Asian J. Biomed. Pharm. Sci. 5.

Ravi, S., Peh, K.K., Darwis, Y., Murthy, B.K., Singh, T.R.R., Mallikarjun, C., 2016. Development and characterization of polymeric microspheres for controlled release protein loaded drug delivery system. Indian J. Pharm. Sci. 70, 303. https://doi.org/10.4103/0250-474X.42978

Reddy, K. R., 2000. Controlled-release, pegylation, liposomal formulations: new mechanisms in the delivery of injectable drugs. Ann. Pharmacother. 34, 915–923.

Ronveaux, C.C., Tomé, D., Raybould, H.E., 2015. Glucagon-Like Peptide 1 Interacts with Ghrelin and Leptin to Regulate Glucose Metabolism and Food Intake through Vagal Afferent Neuron Signaling12. J. Nutr. 145, 672–680. https://doi.org/10.3945/jn.114.206029

Ruan, G., Feng, S.-S., 2003. Preparation and characterization of poly(lactic acid)-poly(ethylene glycol)-poly(lactic acid) (PLA-PEG-PLA) microspheres for controlled release of paclitaxel. Biomaterials 24, 5037–5044.

Rüttimann, E.B., Arnold, M., Hillebrand, J.J., Geary, N., Langhans, W., 2009. Intrameal hepatic portal and intraperitoneal infusions of glucagon-like peptide-1 reduce spontaneous meal size in the rat via different mechanisms. Endocrinology 150, 1174–1181. https://doi.org/10.1210/en.2008-1221

Ryan, G.J., Moniri, N.H., Smiley, D.D.,2013a. Clinical effects of once-weekly exenatide for the treatment of type 2 diabetes mellitus. Am. J. Health-Syst. Pharm. AJHP Off. J. Am. Soc. Health-Syst. Pharm. 70, 1123–1131. https://doi.org/10.2146/ajhp120168

Ryan, G.J., Moniri, N.H., Smiley, D.D.,2013b. Clinical effects of once-weekly exenatide for the treatment of type 2 diabetes mellitus. Am. J. Health-Syst. Pharm. AJHP Off. J. Am. Soc. Health-Syst. Pharm. 70, 1123–1131. https://doi.org/10.2146/ajhp120168

Sjöholm, Å., 2010. Liraglutide Therapy for Type 2 Diabetes: Overcoming Unmet Needs. Pharmaceuticals 3, 764–781. https://doi.org/10.3390/ph3030764

Song, Y.-P., Wang, D.-Y., Wang, X.-L., Lin, L., Wang, Y.-Z., 2011. A method for simultaneously improving the flame retardancy and toughness of PLA. Polym. Adv. Technol. 22, 2295–2301. https://doi.org/10.1002/pat.1760

Talsania, T., Anini, Y., Siu, S., Drucker, D.J., Brubaker, P.L., 2005. Peripheral exendin-4 and peptide YY(3-36) synergistically reduce food intake through different mechanisms in mice. Endocrinology 146, 3748–3756. https://doi.org/10.1210/en.2005-0473

Tang-Christensen, M., Larsen, P.J., Göke, R., Fink-Jensen, A., Jessop, D.S., Møller, M., Sheikh, S.P., 1996. Central administration of GLP-1-(7-36) amide inhibits food and water intake in rats. Am. J. Physiol. 271, R848–856.

Tu, F., Lee, D., 2012. Controlling the Stability and Size of Double-Emulsion-Templated Poly(lactic-co-glycolic) Acid Microcapsules. https://doi.org/10.1021/la301498f

Turton, M.D., O’Shea, D., Gunn, I., Beak, S.A., Edwards, C.M.B., Meeran, K., Choi, S.J., Taylor, G.M., Heath, M.M., Lambert, P.D., Wilding, J.P.H., Smith, D.M., Ghatei, M.A., Herbert, J., Bloom, S.R., 1996. A role for glucagon-like peptide-1 in the central regulation of feeding. Nature 379, 69–72. https://doi.org/10.1038/379069a0

Udenfriend, S., Stein, S., Böhlen, P., Dairman, W., Leimgruber, W., Weigele, M., 1972. Fluorescamine: a reagent for assay of amino acids, peptides, proteins, and primary amines in the picomole range. Science 178, 871–872.

Williams, D.L., Baskin, D.G., Schwartz, M.W., 2009. Evidence that intestinal glucagon-like peptide-1 plays a physiological role in satiety. Endocrinology 150, 1680–1687. https://doi.org/10.1210/en.2008-1045

Xu, X.-J., Ye, Y.-H., Tam, J.P., 2006. Peptides: Biology and Chemistry. Springer Science & Business Media.

Yang, Q., Owusu-Ababio, G., 2000. Biodegradable progesterone microsphere delivery system for osteoporosis therapy. Drug Dev. Ind. Pharm. 26, 61–70.

Yüksel, N., Baykara, T., 1997. Preparation of polymeric microspheres by the solvent evaporation method using sucrose stearate as a droplet stabilizer. J. Microencapsul. 14, 725–733. https://doi.org/10.3109/02652049709006822

Zander, M., Madsbad, S., Madsen, J.L., Holst, J.J., 2002. Effect of 6-week course of glucagon-like peptide 1 on glycaemic control, insulin sensitivity, and beta-cell function in type 2 diabetes: a parallel-group study. Lancet Lond. Engl. 359, 824–830. https://doi.org/10.1016/S0140-6736(02)07952-7

